# Global test of the enemy release hypothesis reveals similar patterns of herbivory across native and non-native plants

**DOI:** 10.1101/2024.10.21.619365

**Authors:** Andrea Galmán, Philip G. Hahn, Brian D. Inouye, Nora Underwood, Yanjie Liu, Susan R. Whitehead, William C. Wetzel

## Abstract

The Enemy Release Hypothesis (ERH) proposes that non-native plants escape their co-evolved herbivores and benefit from reduced herbivory in their introduced ranges. Numerous studies have tested this hypothesis, with conflicting results, but previous studies focus on average levels of herbivory and overlook the substantial within-population variability in herbivory, which may provide unique insights into the ERH. We tested differences in mean herbivory and added a novel approach to the ERH by comparing within-population variability in herbivory between native and non-native plant populations. We include several covariates that might mask an effect of enemy release, including latitude, regional plant richness, plant growth form and plant cover. We use leaf herbivory data collected by the Herbivory Variability Network for 788 plant populations (616 native range populations and 172 introduced range populations) of 503 different native and non-native species distributed worldwide. We found no overall differences in mean herbivory or herbivory variability between native and non-native plant populations. Taken together, our results indicate no evidence of enemy release for non-native plants, suggesting that enemy release is not a generalized mechanism favoring the success of non-native species.

**OPEN RESEARCH STATEMENT:** Data published and available for peer review at https://doi.org/10.5061/dryad.44j0zpckm

## INTRODUCTION

The striking success of some plant species in non-native ranges has attracted the attention of ecological researchers, as it is an important driver of global change, altering ecosystem functioning and destabilizing biotic interactions (Bellard et al., 2022; Vilà et al., 2011). A key hypothesis for the success of invasive plants is the Enemy Release Hypothesis (ERH) (Keane & Crawley, 2002; Williamson, 1996), which posits that non-native plant species can outcompete native species because they are released from the enemy pressure they experience in their native range. An assumption of the ERH is that enemies are less likely to attack novel plants. Empirical studies have tested this hypothesis by comparing mean levels of herbivory between native and non-native species or between populations of the same species in their native and non-native ranges (e.g. Meijer et al., 2015, 2016). Despite the logical appeal of this hypothesis (Enders et al., 2018) and the elegant simplicity of these tests, results have been mixed, with studies finding that non-natives can experience lower, higher, or similar levels of mean herbivory relative to native plants (Colautti et al., 2004; Liu & Stiling, 2006; Meijer et al., 2016). Some authors have thus concluded that enemy release is not important in plant invasions (Agrawal & Kotanen, 2003; Carrillo-Gavilán et al., 2012; Colautti et al., 2004; Ivison et al., 2023), or at least not consistently important, while others have advocated for studies to examine covariates that could be masking or influencing the importance of enemy release (Brian & Catford, 2023; Catford et al., 2022; Chiuffo et al., 2022). Based on recent work showing that plant populations vary not just in mean herbivory but also in intrapopulation variation in leaf damage (Herbivory Variability Network 2023; Wetzel et al., 2023), we propose and test the hypothesis that a key component of enemy release could be differences in the variability of herbivore attack among plant individuals within native and non-native plant populations.

Intrapopulation variability is a critical feature of biological systems, and ecologists are increasingly recognizing the important role it plays in shaping the outcome of competition, consumer-resource interactions, and population dynamics (Benedetti-Cecchi, 2003; Bolnick et al., 2011; Holyoak & Wetzel, 2020; Inouye, 2005; Shoemaker et al., 2020; Violle et al., 2012; Wetzel et al., 2023). For example, theory indicates that factors that increase the variability of herbivore attack among individuals within plant populations should stabilize plant-herbivore interactions and reduce the probability of plant extirpation (Anderson & May, 1978; Bjørnstad & Hansen, 1994; Crawley, 1983). Indeed, when herbivory is aggregated only on a few plant individuals within a population, many plants can escape top-down pressure from herbivores, thus a higher equilibrium population size can be achieved (Anderson & May, 1978; Crawley, 1983). Aggregation also means that herbivores experience greater negative density-dependent feedback which can stabilize the host-herbivore dynamic (Mutz & Inouye, 2023). Thus, for similar levels of mean herbivory, the impact of herbivores might be greater in a population where all plant individuals are attacked than in a population that exhibits high variability in herbivory attack. This suggests that a form of enemy release could occur if herbivore damage were more variable in populations of non-natives than natives, even if mean rates of herbivory across populations were similar between natives and non-natives. Regardless of their potential consequences, differences in intrapopulation variability in herbivory between native and non-native populations would indicate differences in interactions with herbivores.

A key factor likely influencing intrapopulation herbivory variability is how plants are recognized by their potential enemies (Wetzel et al. 2023). We hypothesize that, depending on how host recognition differs between native and non-native plants, non-natives could exhibit higher, lower, or similar levels of intrapopulation variation in herbivory. For example, non-natives would exhibit higher variability in herbivory if most non-native individuals are not recognized by herbivores owing to their novelty but individuals that are recognized (or sampled) by herbivores suffer high damage. High damage on the few unlucky individuals that are used as hosts might be expected because many non-natives are competitive species that prioritize growth over defense (Fahey et al., 2022; Huang et al., 2020; Van Kleunen et al., 2010). Alternatively, non-natives could exhibit lower variability in herbivory than natives because they are less likely to be recognized as hosts by specialist than by generalist herbivores (Goßner et al., 2009; Parker & Hay, 2005; Parker et al., 2006). Specialists often have patchy distributions, potentially leading to more variable damage (Price, 2003) on natives, whereas the generalist-dominated herbivore community on non-natives may leave more homogeneous damage (Joy Massad et al., 2024). Finally, non-native status might be a poor predictor of host recognition, leading to no overall differences in herbivory variability between natives and non-natives. This result would support the perspective that non-native status is a poor predictor of ecological roles relative to functional traits (Agrawal & Kotanen, 2003; Lundgren et al., 2024).

A key recognition from the literature about the ERH over the last decade is that enemy release can vary with geographic and ecological context, both of which can have large influences on species interactions (Brian & Catford, 2023; Catford et al., 2022; Chiuffo et al., 2022; Gioria et al., 2023; Xu et al., 2021). For example, recent studies testing the hypothesis that enemy release varies with latitude for native and non-native plants (i.e., non-parallel latitudinal gradients in herbivory for natives and non-natives) have yielded contrasting results (Allen et al., 2017; Bezemer et al., 2014; Cronin et al., 2015). While some studies focusing on a model species found that enemy release gets weaker with increasing latitude (Bezemer et al., 2014; Guo, 2024), a recent global study reported no correlation between latitude and enemy release (Xu et al., 2021). We argue that there are two important gaps in this literature. First, these studies only examined latitude and not other important factors that influence plant-herbivore interactions, such as plant diversity, growth form or plant cover. Second, ecological context should influence how enemy release affects variability in herbivory as well as mean levels of herbivory. Past studies of geographical and ecological variation in enemy release have only considered mean levels of herbivory and did not examine potential changes in variability in herbivory among individuals, which has been shown to increase with latitude (Herbivory Variability Network, 2023).

The importance of enemy release in shaping the amount and variability of herbivory across plants likely depends on several ecological factors. We identify two mechanisms by which ecological factors might influence enemy release strength. First, enemy release might depend on factors such as latitude and plant diversity, which are predictors of herbivore abundance and richness (Crutsinger et al., 2006; Schemske et al., 2009; Zhang et al., 2016). When considering mean herbivory levels, enemy release might be easier to detect in environments with low herbivore richness (such as low plant diversity environments or temperate regions). However, in environments with high herbivore abundance and richness (such as high plant diversity environments or tropical regions), non-native plants might have a higher risk of being detected and attacked by some herbivore species, leading to high herbivory means for natives and non-natives via an amplification effect (i.e. an increase in species diversity may increase attack risk; Keesing et al., 2006). In environments with high herbivore abundance and richness, we predict that enemy release could be detected by comparing variability in herbivory, with non-natives exhibiting higher variability than natives, because natives should be consistently attacked by their generalist and specialist herbivores while non-natives will either be overlooked or heavily attacked when found (mostly by generalist herbivores, but potentially by some specialist species). Second, enemy release might depend on plant characteristics such as growth form or percent cover that influence how easily a plant is detected by herbivores (Plant Apparency; Feeny, 1976; Galmán et al., 2018; Strauss et al., 2015). Plants with characteristics that make them more obvious hosts for herbivores (woody species or high-cover plants) should have higher levels of attack regardless of native status, but, within a population, non-native plants should have higher variability than natives because natives should be attacked more consistently by a greater number of herbivore species.

Understanding how non-native plants interact with the native herbivores in their introduced range requires work that examines how factors such as latitude, plant diversity, plant growth form or plant cover influence both the mean and variability of herbivory in native and non-native populations. Previous large-scale studies only investigated a limited number of plant species (but see Xu et al 2021) and data for these studies were obtained using different methods, making comparisons across systems difficult (Meijer et al 2016). Here, we perform a global analysis of the Enemy Release Hypothesis using the largest dataset thus far and using a common protocol across all species. We use herbivory data from surveys conducted by the Herbivory Variability Network (https://herbvar.org/). With this study, we present a novel evaluation of the ERH by comparing native and non-native plants both in terms of mean levels of herbivory and within-population variability in the distribution of herbivory. We conduct i) a large biogeographical analysis across 788 populations (616 native and 172 introduced) of 503 plant species; and ii) an analysis comparing native and introduced ranges in a subset of ten species for which we collected survey data from both parts of the range. We test 1) the generality of enemy release by comparing herbivory rates and herbivory variability between native and non-native populations of many species; and 2) whether the effect of enemy release is modulated by ecological context by analyzing the effects of factors influencing herbivore abundance and richness (latitude and plant richness) and factors modulating the interaction between plants and herbivores (plant growth form and focal plant cover).

## MATERIALS AND METHODS

### Field Surveys

This study was conducted using data collected by the Herbivory Variability Network (HerbVar, https://herbvar.org), a team of researchers from 34 countries that aims to better understand the role of variability in the ecology and evolution of plant-herbivore interactions. The dataset includes surveys of 788 plant populations (616 corresponding to plants in their native range and 172 plant populations in their introduced range) encompassing 503 plant species from 135 plant families across 34 countries and six continents (Figure 1).

**Figure 1.**
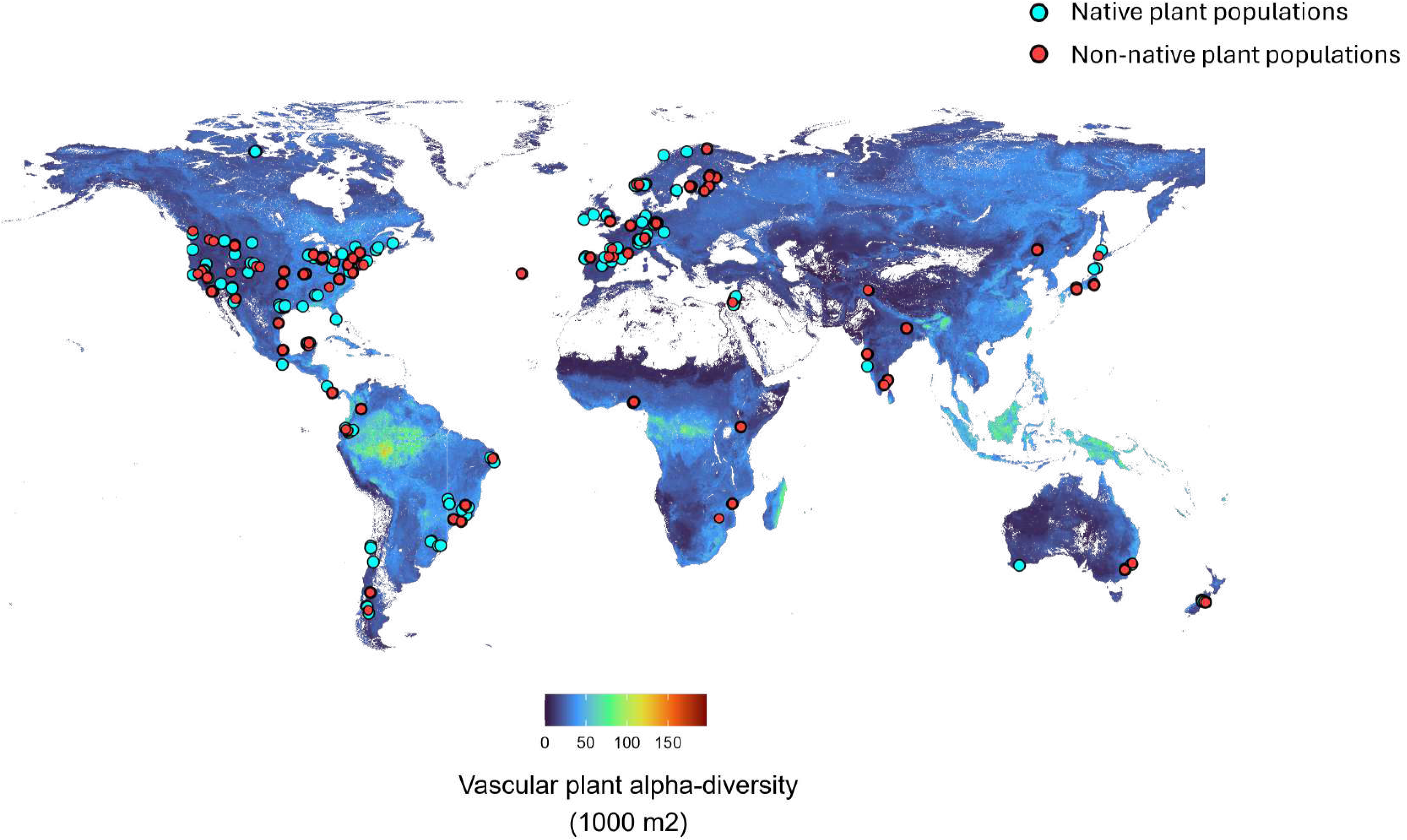
Geographic and environmental locations for the native (purple dots, n=616) and non-native (yellow dots, n = 172) populations of the studied species. The map shows the global distribution of estimated vascular plant alpha diversity (spatial grains 1000 m^2^).

All collaborators followed a standardized protocol (the protocol can be found at the HerbVar website: https://herbvar.org and in the supporting information). In brief, for each survey we randomly chose 30 plant individuals in a population and each of their nearest conspecific neighbors for a total of 60 plant individuals. When the population had less than 90 individuals, we surveyed all the individuals in the population. Our large sample size allows for a robust estimation of variability within a population, as well as mean herbivore damage. For each plant we visually estimated the aboveground proportion of herbivore damage, following a detailed guide. We included invertebrate and vertebrate damage and chewing and mining damage. We examined all above-ground tissues for plants under 2 m tall, while for plants under 2m, we randomly sampled 30 leaves per plant. To estimate the local abundance of the focal species, we also recorded the percent cover of the focal plant species in the sampled area, the sample area for each population was calculated taking into account the density of focal plants, ranging from an area of 0.4 m radius (density between 6 and 10 plants/m^2^) to 3.6 m (for a density ≤ 0.1 plants/m^2^).

### Data acquisition

#### Plant status

We classified each population as native or non-native based on the information provided by the scientific collaborator of each specific region; when the information was not provided, we checked the species status in the Plants of the World Online (POWO, 2024) databases. In addition, we checked the status of the non-native species in the Global Naturalized Alien Flora (GloNAF; van Kleunen et al., 2019). All non-native species in our study (except ten populations of seven non-native plant species) are naturalized and widespread in their introduced ranges (Table S1 in supporting information).

#### Plant diversity data

We extracted plant species richness (estimated number of plant species per 1000m2) for our study sites from sPlotOpen, which predicts plant diversity from a combination of global vegetation surveys and mathematical models (Bruelheide et al., 2019; Sabatini et al., 2022). Finally, note that because sPlotOpen does not provide uncertainty measures for the estimates of plant diversity we were unable to include those in our analyses, and thus all results for plant diversity should be interpreted cautiously.

### Statistical analyses

For our analyses, we use the mean herbivory within a population and variability in herbivory among individuals within a population as separate response variables. For mean herbivory, we averaged the proportion of aboveground herbivory across all individuals surveyed in a population. For each population, we summarize the amount of variability in the proportion of herbivory across individuals by calculating the Gini coefficient using the R package DescTools (Signorell, 2019). The Gini coefficient (range 0-1) represents the level of variation or unevenness of a distribution of a variable among units. The Gini coefficient has certain advantages over other more widely known coefficients of variation; it is calculated with L-moments instead of conventional moments, making it more robust to outliers and more reliable at small sample sizes (Valbuena et al., 2017).

We use Bayesian phylogenetic generalized linear mixed models (GLMM) in R in the brms package (Bürkner, 2021) in R version 4.3.0 (R Core Team, 2023). We used a beta response distribution because it is well suited to represent variables on the 0–1 interval (Douma & Weedon, 2019). Since the beta distribution is undefined for 1, we truncated three values in our data to 0.99. Models ran across seven MCMC chains for at least 5000 total iterations. We assessed runs by ensuring all Rhat values were < 1.03, and visually checked fits via posterior predictive checks. For prior distributions we used normal (0, 2) for slopes, normal (0, 2) for intercepts, gamma (1, 0.05) for phi [the beta distribution dispersion parameter], and cauchy (0, 1) for the standard deviation of random effects. To account for phylogenetic correlations, we built a phylogenetic tree for the species in our study using the phylo.maker function in ‘V.PhyloMaker’ (Jin & Qian, 2019) and ‘ape’ (Paradis & Schliep, 2019) R-packages by matching the family, genus and species epithet from our survey with those in the backbone using the GBOTB.extended phylogeny (i.e., the mega-tree implemented in the ‘V.PhyloMaker’ R package). For each model, we report effect sizes, 95% credible intervals (CIs), Bayes Factors (BF) and marginal Bayesian R2 values.

#### Global differences in herbivory mean and variability between native and non-native populations

We compare herbivory between native and non-native plant populations of different species globally distributed and between populations of the same species in their native and non-native ranges. We ran Bayesian phylogenetic GLMM using mean herbivory and variability in herbivory (Gini) as response variables and plant status (i.e. native or non-native) as a fixed effect. For the global dataset of 772 populations of 503 species, we include plant species and plant phylogeny as random effects. We ran the same models (without the plant phylogeny) for the biogeographical subset of ten species (Table S2 in supporting information) occurring in both native and introduced ranges. For the latter, we also included the interaction with species as a fixed effect.

#### Effect of ecological context modulating Enemy Release

To test whether enemy release was contingent on factors like plant diversity, plant cover, latitude or growth form, we again ran Bayesian phylogenetic GLMM using variability in herbivory (Gini) or mean herbivory as response variables. In this case, for the global dataset of 772 populations of 503 species, we ran different models using plant status as a fixed factor (i.e. native or non-native) and its interaction with i) plant diversity, ii) latitude (absolute values), iii) focal plant cover and iv) growth form (woody versus non-woody species). We included plant species and plant phylogeny as random effects.

For the interactions with plant diversity and plant cover, we ran the same models for the biogeographical subset of ten species occurring in both native and introduced ranges including the interaction with species as a fixed effect.

## RESULTS

### Global differences in herbivory mean and variability between native and non-native populations

Both mean herbivory and variability in herbivory were similar between native and non-native species across the globe. Mean herbivory averaged 5% for both native (95% CI = 2-10%) and non-native species (95% CI = 3-9%, R^2^ = 6%, BF = 0.04, Figure 2a). Similarly, Gini coefficients were 0.60 (0.4-0.8) for native species and 0.62 (0.4-0.8) for non-native species (R^2^ = 5%, BF = 0.09, Figure 2b).

**Figure 2:**
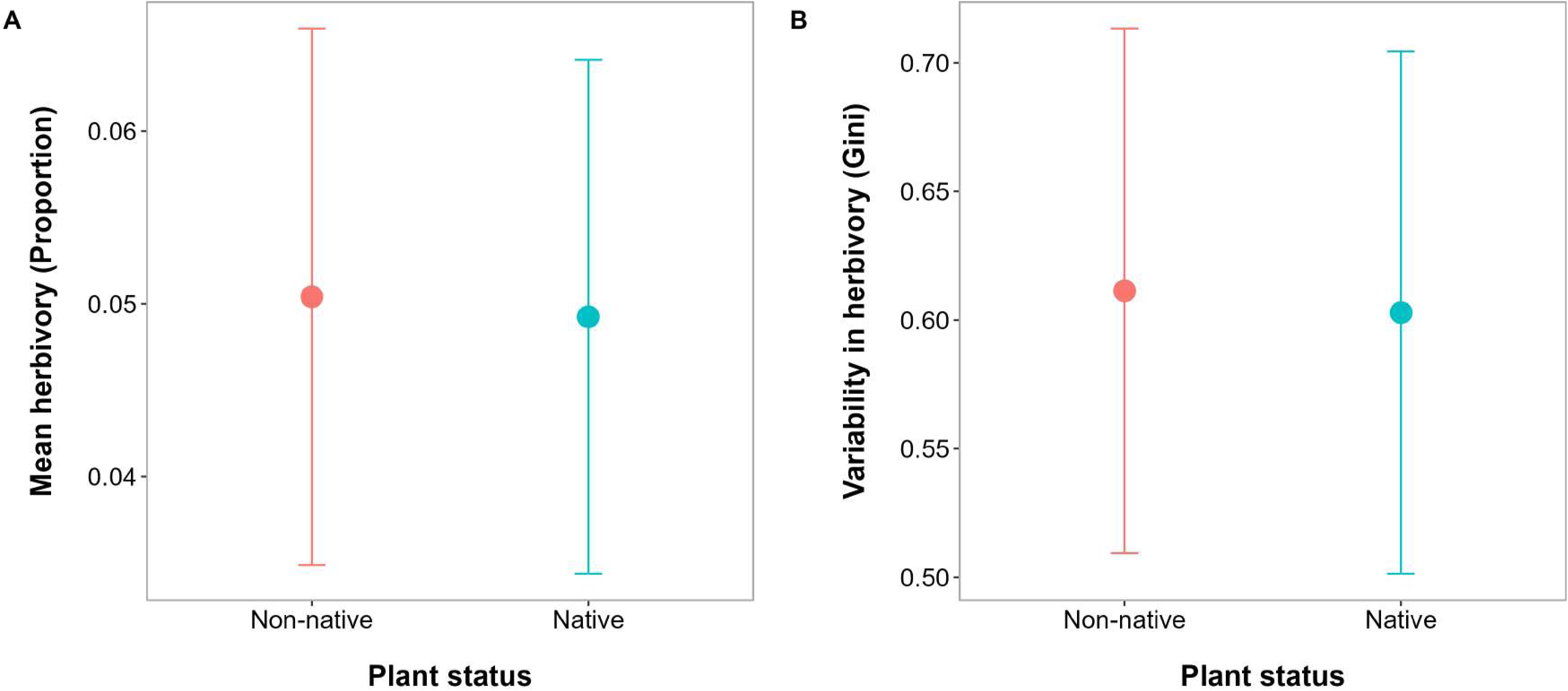
Results from the global analysis of enemy release showing no differences in (a) mean herbivory and (b) variability in herbivory (Gini coefficient) between native (blue) and non-native (red) species. Dots show predicted means and lines 95% credible intervals from Bayesian phylogenetic beta regressions.

Non-native status also had no effect on herbivory patterns within species when we restricted our dataset to the ten species with surveys in both their native and introduced ranges. Mean herbivory and herbivory variability were similar between native and non-native populations within plant species (supporting information, Figures S1 and S2).

### Effect of ecological context modulating Enemy Release: latitude

Mean herbivory decreased with increasing latitude from 8% (95% CI: 4-15 %) at the equator to 3% (95% CI: 2-5%) at 70° N/S (R2 = 5%, BF = 3.7, Figure 3). In contrast, variability in herbivory increased with increasing latitude from Gini=0.5 (95% CI: 0.3-0.7) at the equator to Gini= 0.7 (95% CI: 0.5-0.9) at 70° N/S (R2 = 5%, BF = 1.2, Figure 3). The data did not support an interaction between non-native status and latitude for mean damage (Estimate = 0.01, 95% CI=0-0.02, R2 = 6 %, BF = 0.08, Figure 3c) or for variability in herbivory (Estimate = 0.5, 95% CI = 0.43-0.57, R2 = 5%, BF = 0.07, Figure 3d).

**Figure 3:**
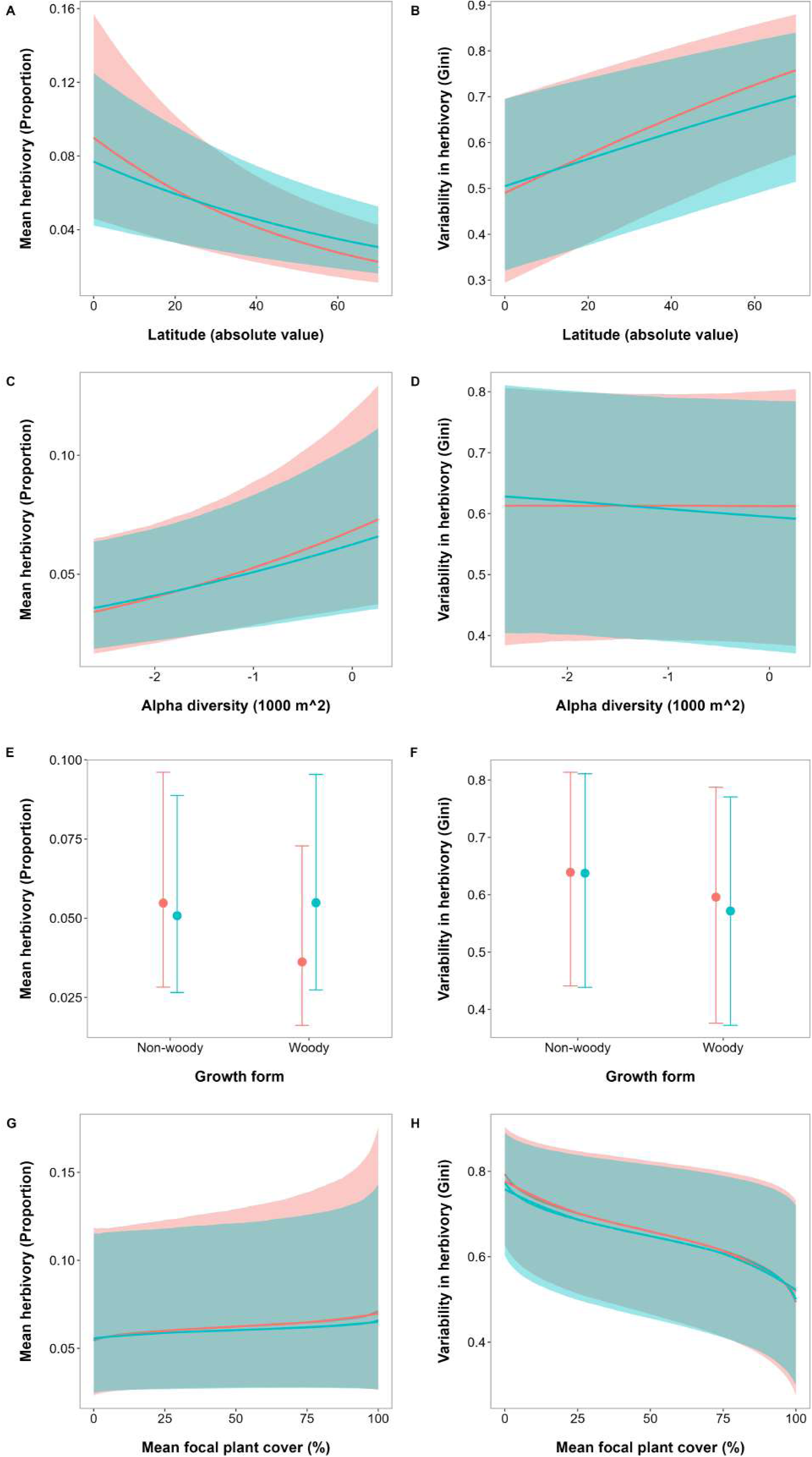
Results from the analyses of ecological factors potentially driving differences in herbivory for native (blue) and non-native (red) species populations. There were no differences in mean herbivory or herbivory variability between native and non-native populations at different latitudes (a, b), different levels of plant alpha diversity (c, d), nor considering different growth forms (e, f) or with different focal plant cover (g, h). Graphs show predicted means and 95% credible intervals from Bayesian phylogenetic beta regressions.

### Effect of ecological context modulating Enemy Release: plant diversity

Mean herbivory increased with plant diversity; mean herbivory increased from 4% (95% CI: 2-7%) at the lower levels of diversity (7 plant species per 1000m^2^) to 11% (95% CI: 2-40%) at the greatest levels (57 plant species per 1000m2) (R2 = 4%, BF = 0.8, Figure 3). However, variability in herbivory was not related to plant diversity levels (Gini=0.5, 95% CI= 0.39-0.56, R2 =6 %, BF = 0.08, Figure 3). The data did not support an interaction between non-native status and diversity for mean damage (Estimate = 0.5, 95% CI= 0.41-0.56, R2 = 6 %, BF = 0.08, Figure 3c) or for variability in herbivory (Estimate = 0.5, 95% CI= 0.43-0.57, R2 = 6 %, BF = 0.07, Figure 3d), suggesting that plant diversity levels do not determine differences in herbivory between native and non-native plants. When comparing native and introduced ranges for the subset of ten species, we again did not find a relationship between plant diversity and patterns of mean herbivory or variability in herbivory between the native and introduced range of each species (supporting information, Figure S3 and S4).

### Effect of ecological context modulating Enemy Release: plant growth form

There was no effect of growth from in herbivory mean (Estimate = −0.4, 95% CI= −0.99-0.09, R2 = 5%, BF = 0.45, Figure 3E) or herbivory variability (Estimate = −0.18, 95% CI= −0.62-0.24, R2 = 5%, BF = 0.15, Figure 3F). In addition, we found no effect of growth form (i.e. woody versus non-woody species) determining differences in mean herbivory (Estimate = 0.51, 95% CI= −0.04-1.08, R2 = 5%, BF = 0.75, Figure 3e) or herbivory variability (Estimate = −0.09, 95% CI= −0.56-0.37, R2 = 5%, BF = 0.12, Figure 3f) between native and non-native species.

### Effect of ecological context modulating Enemy Release: focal plant cover

There was no effect of focal plant cover in herbivory mean (Estimate = 0.04, 95% CI= −0.99-0.17, R2 = 4 %, BF = 0.04, Figure 3G) but we found a negative effect of focal plant cover on variability in herbivory (Estimate = −0.19, 95% CI= [-0.29, −0.08], R2 =5%, BF =14.42, Figure 3h). However, there was no effect of the interaction between focal plant cover and non-native status for mean herbivory (Estimate = −0.01, 95% CI = −0.17, 0.14, R2 =4%, pp =1, BF = 0.04, Figure 3g) or herbivory variability (Estimate = 0.02, 95% CI = −0.11, 0.15, R2 = 5%, pp =1, BF = 0.03, Figure 3h), suggesting that plant cover influences herbivory in a similar way for both native and non-native plants. When comparing native and introduced ranges for the subset of ten species, we did not find an effect of cover determining differences in mean or variability in herbivory for any of the species (supporting information, Figure S5 and S6).

## DISCUSSION

Using a global survey of herbivore damage on 503 plant species collected with a standardized protocol, we evaluated a main prediction of the Enemy Release Hypothesis, that herbivory is lower on non-native than native plants. In addition, we proposed an expansion of the ERH to include potential differences in the variability of herbivory. Embracing variability as a key ecological response variable can broaden understanding of ecological processes, including plant-herbivore interactions (Wetzel et al. 2023, Herbivory Variability Network 2023). We predicted that variability could differ between native and non-native plants if, based on a lack of long-term coevolutionary history between non-native species and the native herbivores, herbivore host-recognition differed between native and non-native plant species. We compared the mean and variability of herbivore leaf damage between native and non-native plant populations and analyzed the effect of biotic and abiotic factors potentially masking enemy release. Despite this expanded perspective on enemy release and despite the breadth and intensity of our sampling, we found no differences in either mean damage or variability in herbivory between natives and non-natives, suggesting that there are no overall differences in plant-herbivore interactions between native and non-native plants. Below we discuss potential explanations for why we observed no differences in herbivory patterns between native and non-natives and the implications of this finding for invasion biology.

A previously proposed explanation for findings of no difference in herbivory between natives and non-natives is that enemy release is apparent only after accounting for key ecological covariates that influence herbivory and could mask enemy release (Brian & Catford, 2023; Catford et al., 2022; Chiuffo et al., 2022; Gioria et al. 2023). However, after we accounted for variation in herbivory with latitude and plant diversity, growth form, and abundance—factors that are key determinants of herbivory—we still found no differences between natives and non-natives. It is not that these factors had no impact on herbivory on non-natives. To the contrary, latitude, plant community diversity, and plant abundance had strong relationships with both the mean and variability of herbivory on non-natives. These relationships, however, were strikingly similar to those for natives. A key implication of this result is that, on average, non-natives might be more ecologically similar to natives in plant-herbivore interactions than is often hypothesized. For example, the similarity in the relationship between local abundance (percent cover) and herbivory variability for natives and non-natives suggests that ecological factors like abundance are more important for herbivore host recognition and use than the distinction between native and non-native. Low abundance, locally rare populations may experience high variability in damage—rare individuals either escape notice or are found and highly consumed— regardless of whether they have native or non-native status. More generally, these results suggest that the broad environmental and ecological factors we examined are more important determinants of herbivory patterns than non-native status.

One explanation for the lack of differences in herbivory patterns between native and non-native plants is that enemy release is not determined by whether a plant is native or not but by the specific characteristics of each plant species. The interaction of non-native plants with native herbivores may be determined by the functional traits of plants and by the functional similarity between non-native plant species and the native community. Accordingly, previous studies showed that phylogenetic relatedness and trait similarity to native species predict herbivory in non-native species (Pearse & Hipp, 2009; Pearse & Rosenheim, 2020). This notion is consistent with recent studies on invasion ecology that highlight the predominant role of functional traits for the success of non-native species (El-Barougy et al., 2020; Qian & Sandel, 2022), indicating that non-native species functionally similar to the natives benefit from preadaptation to the novel environment and are more likely to naturalize. One implication of the fact that interactions with native herbivores depend on the non-native traits is that enemy release is not widespread. More likely, only a fraction of non-native species benefit from enemy release when introduced into a novel environment due to differences in host detection, depending on the environment and the specific traits of the species. Indeed, we hypothesize that enemy release may be important only for those rare non-native species that differ significantly in traits from their native competitors.

Another explanation for the lack of differences in herbivory between native and non-native plants is that enemy release, though potentially an important factor in the early stages of plant invasions, loses its importance once non-native populations are established, as is the case for the non-native species in our study. A waning of enemy release would make sense because studies show that native herbivores adapt to use non-native hosts as a resource over time, often surprisingly quickly (Ivison et al., 2023; Parker & Gilbert 2004; Mitchell et al. 2010). Future studies could test this hypothesis by selecting species in the early stages of invasions and comparing herbivory or even experimentally excluding or adding herbivores. Regardless, our data provide a robust test of the strength of enemy release for established non-natives across a broad sampling of geography, growth form, and taxonomy.

Overall, our results indicate that not all non-native species escape herbivory in their exotic ranges, as a consequence enemy release is not an overall mechanism favouring invasion success. Non-native species may present competitive advantages to natives based on different mechanisms than escaping their enemies. For instance, the success of non-native species is not only determined by how resistant they are to the native herbivores in the new environments but also by how tolerant they are to herbivore attacks. In this regard, invasive plants have been found to perform better than natives under similar damage levels (Ashton & Lerdau, 2008), suggesting that even when non-native species do not benefit from enemy release, they might present different mechanisms than natives to tolerate or overcompensate afterattack (Liao et al., 2014) which could help them outcompete the native plants in the community. Alternatively, other studies have suggested that arbuscular mycorrhizal fungi and soil nitrogen levels may be critical in mediating the promotion of introduced plants (Zhang et al., 2023).

Finally, the fact that herbivore damage is not generally lower for non-natives than natives suggests that introducing herbivores for biocontrol will not be a silver bullet for managing established non-native species. Indeed, our results are consistent with the observation that controlling non-native plants via classical biocontrol is challenging (Shen et al., 2023). Management strategies must be specific to the herbivore community and the functional characteristics of the non-native species. Managers should not assume that theory can guide their decisions. Instead, they will need to use experimental ecology methods to elucidate the factors in their systems that determine the success of their non-native species and should include measures of performance or tolerance after damage to fully understand the impacts of trophic interactions on introduced species. In cases where herbivores are used as management tools, our results suggest that successful non-native management via enemies could be thought of as increasing herbivory on non-natives above what is natural rather than restoring enemy pressure that may not have been escaped in the first place.

## REFERENCES

Agrawal, A. A., & Kotanen, P. M. (2003). Herbivores and the success of exotic plants: a phylogenetically controlled experiment. Ecology Letters, 6(8), 712–715. 10.1046/j.1461-0248.2003.00498.x

Agrawal, A. A., Lau, J. A., & Hambäck, P. A. (2006). Community heterogeneity and the evolution of interactions between plants and insect herbivores. The Quarterly Review of Biology, 81(4), 349–376. 10.1086/511529

Allen, W. J., Meyerson, L. A., Cummings, D., Anderson, J., Bhattarai, G. P., & Cronin, J. T. (2017). Biogeography of a plant invasion: drivers of latitudinal variation in enemy release. Global Ecology and Biogeography, 26(4), 435–446. 10.1111/geb.12550

Anderson, R. M., & May, R. M. (1978). Regulation and Stability of Host-Parasite Population Interactions: I. Regulatory Processes. The Journal of Animal Ecology, 47(1), 219. 10.2307/3933

Bellard, C., Marino, C., & Courchamp, F. (2022). Ranking threats to biodiversity and why it doesn’t matter. Nature Communications, 13(1), 1–4.

Benedetti-Cecchi, L. (2003). The importance of the variance around the mean effect size of ecological processes. Ecology, 84(9), 2335–2346. 10.1890/02-8011

Bezemer, T. M., Harvey, J. A., & Cronin, J. T. (2014). Response of Native Insect Communities to Invasive Plants. Annual Review of Entomology, 59(1), 119–141. 10.1146/annurev-ento-011613-162104

Bjørnstad, O. N., & Hansen, T. F. (1994). Individual variation and population dynamics. Oikos, 69(1), 167–171.

Bolnick, D. I., Amarasekare, P., Araújo, M. S., Bürger, R., Levine, J. M., Novak, M., Rudolf, V. H. W., Schreiber, S. J., Urban, M. C., & Vasseur, D. A. (2011). Why intraspecific trait variation matters in community ecology. Trends in Ecology & Evolution, 26(4), 183–192. 10.1016/j.tree.2011.01.009

Brian, J. I., & Catford, J. A. (2023). A mechanistic framework of enemy release. Ecology Letters, 26(12), 2147–2166. 10.1111/ele.14329

Bruelheide, H., Dengler, J., Jiménez-Alfaro, B., Purschke, O., Hennekens, S. M., Chytrý, M., Pillar, V. D., Jansen, F., Kattge, J., Sandel, B., Aubin, I., Biurrun, I., Field, R., Haider, S., Jandt, U., Lenoir, J., Peet, R. K., Peyre, G., Sabatini, F. M., … Zverev, A. (2019). sPlot – A new tool for global vegetation analyses. Journal of Vegetation Science, 30(2), 161–186. 10.1111/jvs.12710

Bürkner, P.-C. (2021). Bayesian Item Response Modeling in R with brms and Stan. Journal of Statistical Software, 100(5). 10.18637/jss.v100.i05

Carrillo-Gavilán, A., Moreira, X., Zas, R., Vilà, M., & Sampedro, L. (2012). Early resistance of alien and native pines against two native generalist insect herbivores: no support for the natural enemy hypothesis. Functional Ecology, 26(1), 283–293. 10.1111/j.1365-2435.2011.01931.x

Catford, J. A., Wilson, J. R. U., Pyšek, P., Hulme, P. E., & Duncan, R. P. (2022). Addressing context dependence in ecology. Trends in Ecology & Evolution, 37(2), 158–170. 10.1016/j.tree.2021.09.007

Chiuffo, M. C., Moyano, J., Policelli, N., Torres, A., Vitali, A., Nuñez, M. A., & Rodriguez-Cabal, M. A. (2022). Importance of invasion mechanisms varies with abiotic context and plant invader growth form. Journal of Ecology, 110(8), 1957–1969. 10.1111/1365-2745.13929

Colautti, R. I., Ricciardi, A., Grigorovich, I. A., & MacIsaac, H. J. (2004). Is invasion success explained by the enemy release hypothesis? Ecology Letters, 7(8), 721–733. 10.1111/j.1461-0248.2004.00616.x

Crawley, M. J. (1983). Herbivory: The Dynamics of Animal-Plant Interactions. (pp. 270–275). University of California Press.

Cronin, J. T., Bhattarai, G. P., Allen, W. J., & Meyerson, L. A. (2015). Biogeography of a plant invasion: plant–herbivore interactions. Ecology, 96(4), 1115–1127. 10.1890/14-1091.1

Crutsinger, G. M., Collins, M. D., Fordyce, J. A., Gompert, Z., Nice, C. C., & Sanders, N. J. (2006). Plant Genotypic Diversity Predicts Community Structure and Governs an Ecosystem Process. Science, 313(5789), 966–968. 10.1126/science.1128326

Douma, J. C., & Weedon, J. T. (2019). Analysing continuous proportions in ecology and evolution: A practical introduction to beta and Dirichlet regression. Methods in Ecology and Evolution, 10(9), 1412–1430. 10.1111/2041-210X.13234

El-Barougy, R. F., Elgamal, I., Rohr, R. P., Probert, A. F., Khedr, A. A., & Bacher, S. (2020). Functional similarity and dissimilarity facilitate alien plant invasiveness along biotic and abiotic gradients in an arid protected area. Biological Invasions, 22(6), 1997–2016. 10.1007/s10530-020-02235-3

Enders, M., Hütt, M., & Jeschke, J. M. (2018). Drawing a map of invasion biology based on a network of hypotheses. Ecosphere, 9(3). 10.1002/ecs2.2146

Fahey, C., Koyama, A., & Antunes, P. M. (2022). Vulnerability of non-native invasive plants to novel pathogen attack: do plant traits matter? Biological Invasions, 24(11), 3349–3379. 10.1007/s10530-022-02853-z

Feeny, P. (1976). Plant Apparency and Chemical Defense. In Biochemical Interaction Between Plants and Insects (pp. 1–40). Springer US. 10.1007/978-1-4684-2646-5_1

Galmán, A., Abdala-Roberts, L., Zhang, S., Berny-Mier y Teran, J. C., Rasmann, S., & Moreira, X. (2018). A global analysis of elevational gradients in leaf herbivory and its underlying drivers: Effects of plant growth form, leaf habit and climatic correlates. Journal of Ecology, 106(1), 413–421. 10.1111/1365-2745.12866

Gioria, M., Hulme, P. E., Richardson, D. M., & Pyšek, P. (2023). Why Are Invasive Plants Successful? Annual Review of Plant Biology, 74(1), 635–670. 10.1146/annurev-arplant-070522-071021

Goßner, M. M., Chao, A., Bailey, R. I., & Prinzing, A. (2009). Native Fauna on Exotic Trees: Phylogenetic Conservatism and Geographic Contingency in Two Lineages of Phytophages on Two Lineages of Trees. The American Naturalist, 173(5), 599–614. 10.1086/597603

Guo, Y., Parepa, M., Wang, H., Wang, M., Wu, J., Li, B., … & Bossdorf, O. (2024). Global heterogeneity of latitudinal patterns in herbivory between native and exotic plants. bioRxiv, 2024–01.

Herbivory Variability Network, Robinson, M. L., Hahn, P. G., Inouye, B. D., Underwood, N., Whitehead, S. R., Abbott, K. C., Bruna, E. M., Cacho, N. I., Dyer, L. A., Abdala-Roberts, L., Allen, W. J., Andrade, J. F., Angulo, D. F., Anjos, D., Anstett, D. N., Bagchi, R., Bagchi, S., Barbosa, M., Barrett, S., … Wetzel, W. C. (2023). Plant size, latitude, and phylogeny explain within-population variability in herbivory. Science, 382(6671), 679–683. 10.1126/science.adh8830

Holyoak, M., & Wetzel, W. C. (2020). Variance-Explicit Ecology: A Call for Holistic Study of the Consequences of Variability at Multiple Scales. In Unsolved Problems in Ecology (pp. 25–42). Princeton University Press. 10.1515/9780691195322-005

Huang, K., Kong, D.-L., Lu, X.-R., Feng, W.-W., Liu, M.-C., & Feng, Y.-L. (2020). Lesser leaf herbivore damage and structural defense and greater nutrient concentrations for invasive alien plants: Evidence from 47 pairs of invasive and non-invasive plants. Science of The Total Environment, 723, 137829. 10.1016/j.scitotenv.2020.137829

Inouye, B. D. (2005). The Importance of the Variance around the Mean Effect Size of Ecological Processes: Comment. Ecology, 86(1), 262–265. http://www.jstor.org/stable/3451006

Ivison, K., Speed, J. D. M., Prestø, T., & Dawson, W. (2023). Testing enemy release of non-native plants across time and space using herbarium specimens in Norway. Journal of Ecology, 111(2), 300–313. 10.1111/1365-2745.13998

Jin, Y., & Qian, H. (2019). V.PhyloMaker: an R package that can generate very large phylogenies for vascular plants. Ecography, 42(8), 1353–1359. 10.1111/ecog.04434

Johnson, M. T. J., & Rasmann, S. (2011). The latitudinal herbivory-defence hypothesis takes a detour on the map. New Phytologist, 191(3), 589–592. 10.1111/j.1469-8137.2011.03816.x

Joy Massad, T., Rangel Nascimento, A., Fernando Campos Moreno, D., Simbaña, W., Garcia Lopez, H., Sulca, L., Lepesqueur, C., Richards, L. A., Forister, M. L., Stireman, J. O., Tepe, E. J., Uckele, K. A., Braga, L., Walla, T. R., Smilanich, A. M., Grele, A., & Dyer, L. A. (2024). Variation in the strength of local and regional determinants of herbivory across the Neotropics. Oikos, 2024(2). 10.1111/oik.10218

Keane, R. M., & Crawley, M. J. (2002). Exotic plant invasions and the enemy release hypothesis. Trends in Ecology & Evolution, 17(4), 164–170.

Keesing, F., Holt, R. D., & Ostfeld, R. S. (2006). Effects of species diversity on disease risk. Ecology Letters, 9(4), 485–498. 10.1111/j.1461-0248.2006.00885.x

Liu, H., & Stiling, P. (2006). Testing the enemy release hypothesis: a review and meta-analysis. Biological Invasions, 8(7), 1535–1545. 10.1007/s10530-005-5845-y

Liao, Z.-Y., Zheng, Y.-L., Lei, Y.-B., & Feng, Y.-L. (2014). Evolutionary increases in defense during a biological invasion. Oecologia, 174(4), 1205–1214. 10.1007/s00442-013-2852-z

Lundgren, E. J., Bergman, J., Trepel, J., le Roux, E., Monsarrat, S., Kristensen, J. A., Pedersen, R. Ø., Pereyra, P., Tietje, M., & Svenning, J.-C. (2024). Functional traits—not nativeness—shape the effects of large mammalian herbivores on plant communities. Science, 383(6682), 531– 537. 10.1126/science.adh2616

Meijer, K., Schilthuizen, M., Beukeboom, L., & Smit, C. (2016). A review and meta-analysis of the enemy release hypothesis in plant–herbivorous insect systems. PeerJ, 4, e2778. 10.7717/peerj.2778

Meijer, K., Zemel, H., Chiba, S., Smit, C., Beukeboom, L. W., & Schilthuizen, M. (2015). Phytophagous Insects on Native and Non-Native Host Plants: Combining the Community Approach and the Biogeographical Approach. PLOS ONE, 10(5), e0125607. 10.1371/journal.pone.0125607

Moles, A. T., Bonser, S. P., Poore, A. G. B., Wallis, I. R., & Foley, W. J. (2011). Assessing the evidence for latitudinal gradients in plant defence and herbivory. Functional Ecology, 25(2), 380–388. 10.1111/j.1365-2435.2010.01814.x

Mutz, J. and B. D. Inouye (2023). Spatial and ontogenetic variance in local densities modify selection on demographic traits. Functional Ecology 37(6): 1628–1641.

Paradis, E., & Schliep, K. (2019). ape 5.0: an environment for modern phylogenetics and evolutionary analyses in R. Bioinformatics, 35(3), 526–528. 10.1093/bioinformatics/bty633

Parker, J. D., & Hay, M. E. (2005). Biotic resistance to plant invasions? Native herbivores prefer non-native plants. Ecology Letters, 8(9), 959–967. 10.1111/j.1461-0248.2005.00799.x

Parker, J. D., Burkepile, D. E., & Hay, M. E. (2006). Opposing Effects of Native and Exotic Herbivores on Plant Invasions. Science, 311(5766), 1459–1461. 10.1126/science.1121407

Pearse, I. S., & Hipp, A. L. (2009). Phylogenetic and trait similarity to a native species predict herbivory on non-native oaks. Proceedings of the National Academy of Sciences, 106(43), 18097–18102. 10.1073/pnas.0904867106

Pearse, I. S., & Rosenheim, J. A. (2020). Phylogenetic escape from pests reduces pesticides on some crop plants. Proceedings of the National Academy of Sciences, 117(43), 26849–26853. 10.1073/pnas.2013751117

Price, P. W. (2003). Macroevolutionary theory on macroecological patterns. Cambridge University Press.

POWO (2024). “Plants of the World Online. Facilitated by the Royal Botanic Gardens, Kew. Published on the Internet; http://www.plantsoftheworldonline.org/ Retrieved 30 January 2024.” In “Plants of the World Online. Facilitated by the Royal Botanic Gardens, Kew. Published on the Internet; http://www.plantsoftheworldonline.org/ Retrieved 30 January 2024.”

Qian, H., & Sandel, B. (2017). Phylogenetic relatedness of native and exotic plants along climate gradients in California, <scp>USA</scp>. Diversity and Distributions, 23(11), 1323–1333. 10.1111/ddi.12620

Qian, H., & Sandel, B. (2022). Darwin’s preadaptation hypothesis and the phylogenetic structure of native and alien regional plant assemblages across North America. Global Ecology and Biogeography, 31(3), 531–545. 10.1111/geb.13445

R Core Team. (2023). _R: A Language and Environment for Statistical Computing_. R Foundation for Statistical Computing, Vienna, Austria. <https://www.R-project.org/>.

Root, R. B. (1973). Organization of a Plant-Arthropod Association in Simple and Diverse Habitats: The Fauna of Collards (Brassica Oleracea). Ecological Monographs, 43(1), 95–124. 10.2307/1942161

Sabatini, F. M., Jiménez-Alfaro, B., Jandt, U., Chytrý, M., Field, R., Kessler, M., Lenoir, J., Schrodt, F., Wiser, S. K., Arfin Khan, M. A. S., Attorre, F., Cayuela, L., De Sanctis, M., Dengler, J., Haider, S., Hatim, M. Z., Indreica, A., Jansen, F., Pauchard, A., … Bruelheide, H. (2022).Global patterns of vascular plant alpha diversity. Nature Communications, 13(1), 4683. 10.1038/s41467-022-32063-z

Schemske, D. W., Mittelbach, G. G., Cornell, H. V., Sobel, J. M., & Roy, K. (2009). Is There a Latitudinal Gradient in the Importance of Biotic Interactions? Annual Review of Ecology, Evolution, and Systematics, 40(1), 245–269. 10.1146/annurev.ecolsys.39.110707.173430

Shen, C., Chen, P., Zhang, K., He, M., Wan, J., Wang, Y., … & Siemann, E. (2023). Dynamics and mechanisms of secondary invasion following biological control of an invasive plant. New Phytologist, 238(6), 2594–2606.

Signorell, A., Aho, K., Alfons, A., Anderegg, N., Aragon, T., Arppe, A., … & Borchers, H. W. (2019). DescTools: Tools for descriptive statistics. R package version 0.99, 28, 17.

Shoemaker, L. G., Barner, A. K., Bittleston, L. S., & Teufel, A. I. (2020). Quantifying the relative importance of variation in predation and the environment for species coexistence. Ecology Letters, 23(6), 939–950. 10.1111/ele.13482

Stephens, A. E. A., & Myers, J. H. (2012). Resource concentration by insects and implications for plant populations. Journal of Ecology, 100(4), 923–931. 10.1111/j.1365-2745.2012.01971.x

Strauss, S. Y., Cacho, N. I., Schwartz, M. W., Schwartz, A. C., & Burns, K. C. (2015). Apparency revisited. Entomologia Experimentalis et Applicata, 157(1), 74–85. 10.1111/eea.12347

Valbuena, R., Maltamo, M., Mehtätalo, L., & Packalen, P. (2017). Key structural features of Boreal forests may be detected directly using L-moments from airborne lidar data. Remote Sensing of Environment, 194, 437–446. 10.1016/j.rse.2016.10.024

van Kleunen, M., Pyšek, P., Dawson, W., Essl, F., Kreft, H., Pergl, J., Weigelt, P., Stein, A., Dullinger, S., König, C., Lenzner, B., Maurel, N., Moser, D., Seebens, H., Kartesz, J., Nishino, M., Aleksanyan, A., Ansong, M., Antonova, L. A., … Winter, M. (2019). The Global Naturalized Alien Flora (GloNAF) database. Ecology, 100(1). 10.1002/ecy.2542

van Kleunen, M., Weber, E., & Fischer, M. (2010). A meta-analysis of trait differences between invasive and non-invasive plant species. Ecology Letters, 13(2), 235–245. 10.1111/j.1461-0248.2009.01418.x

Vilà, M., Espinar, J. L., Hejda, M., Hulme, P. E., Jarošík, V., Maron, J. L., Pergl, J., Schaffner, U., Sun, Y., & Pyšek, P. (2011). Ecological impacts of invasive alien plants: a meta-analysis of their effects on species, communities and ecosystems. Ecology Letters, 14(7), 702–708.

Violle, C., Enquist, B. J., McGill, B. J., Jiang, L., Albert, C. H., Hulshof, C., Jung, V., & Messier, J. (2012). The return of the variance: intraspecific variability in community ecology. Trends in Ecology & Evolution, 27(4), 244–252. 10.1016/j.tree.2011.11.014

Wetzel, W. C., Inouye, B. D., Hahn, P. G., Whitehead, S. R., & Underwood, N. (2023). Variability in Plant–Herbivore Interactions. Annual Review of Ecology, Evolution, and Systematics, 54(1), 451–474. 10.1146/annurev-ecolsys-102221-045015

Williamson, M. (1996). Biological invasions. Springer Science & Business Media.

Xu, M., Mu, X., Zhang, S., Dick, J. T. A., Zhu, B., Gu, D., Yang, Y., Luo, D., & Hu, Y. (2021). A global analysis of enemy release and its variation with latitude. Global Ecology and Biogeography, 30(1), 277–288. 10.1111/geb.13229

Zhang, K., Lin, S., Ji, Y., Yang, C., Wang, X., Yang, C., Wang, H., Jiang, H., Harrison, R. D., & Yu, D. W. (2016). Plant diversity accurately predicts insect diversity in two tropical landscapes. Molecular Ecology, 25(17), 4407–4419. 10.1111/mec.13770

Zhang, X., Zhang, T., & Liu, Y. (2023). Effects of arbuscular mycorrhizal fungi on plant invasion success driven by nitrogen fluctuations. Journal of Applied Ecology, 60(11), 2425–2436.

Zvereva, E. L., Zverev, V., & Kozlov, M. V. (2022). Insect herbivory increases from forest to alpine tundra in Arctic mountains. Ecology and Evolution, 12(1). 10.1002/ece3.8537

