## supporting information for "Global test of the enemy release hypothesis reveals similar patterns of herbivory across native and non-native plants"

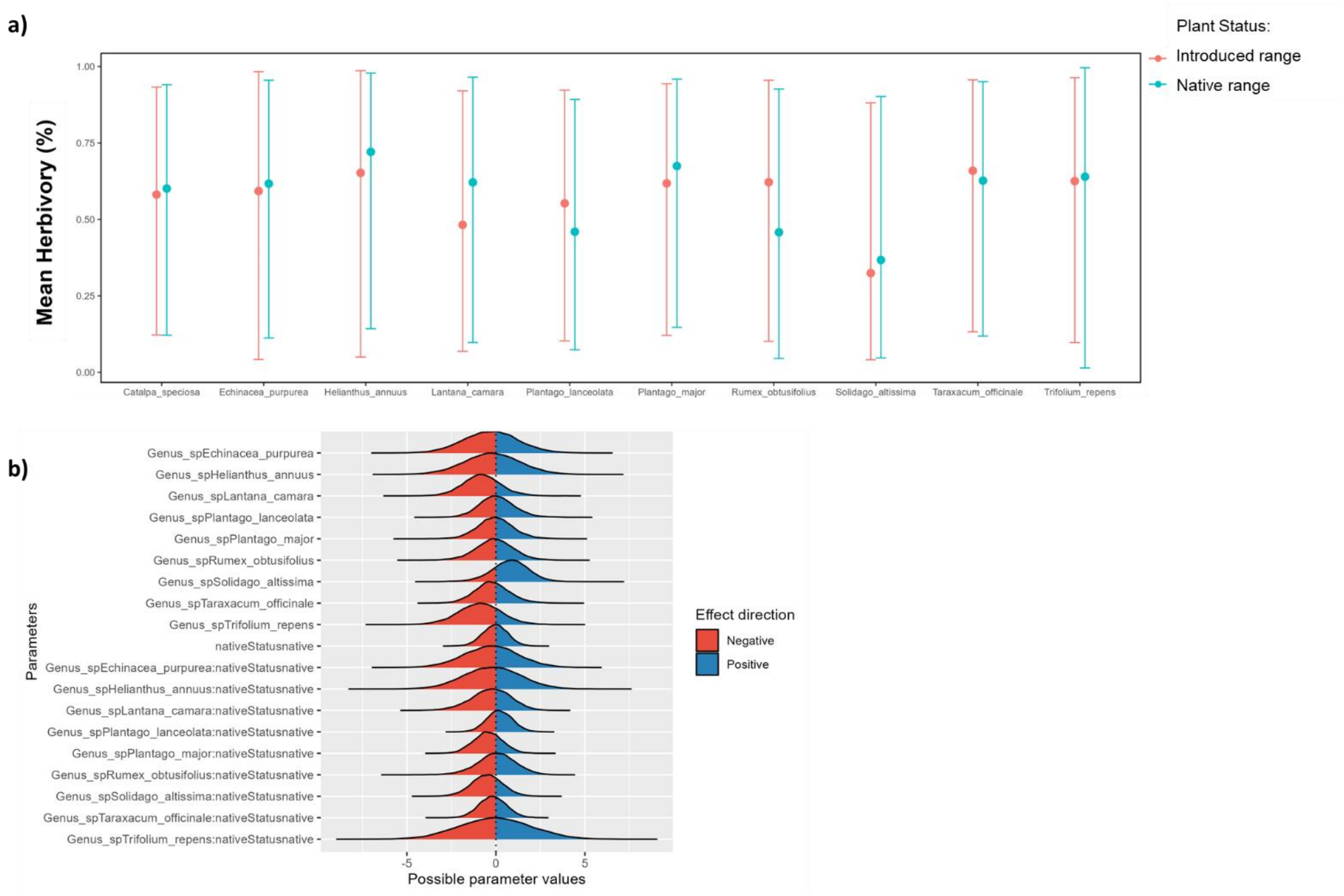


**Figure S1:** Results from models for the biogeographical subsets of ten species for mean herbivory. a) Comparison of mean intrapopulation variability in herbivory for populations in their introduced (in red) and native (in blue) range for the subset of 10 species included in the biogeographic analysis and b) evaluation plot with the proportion of the posterior distribution of the factors in the model.

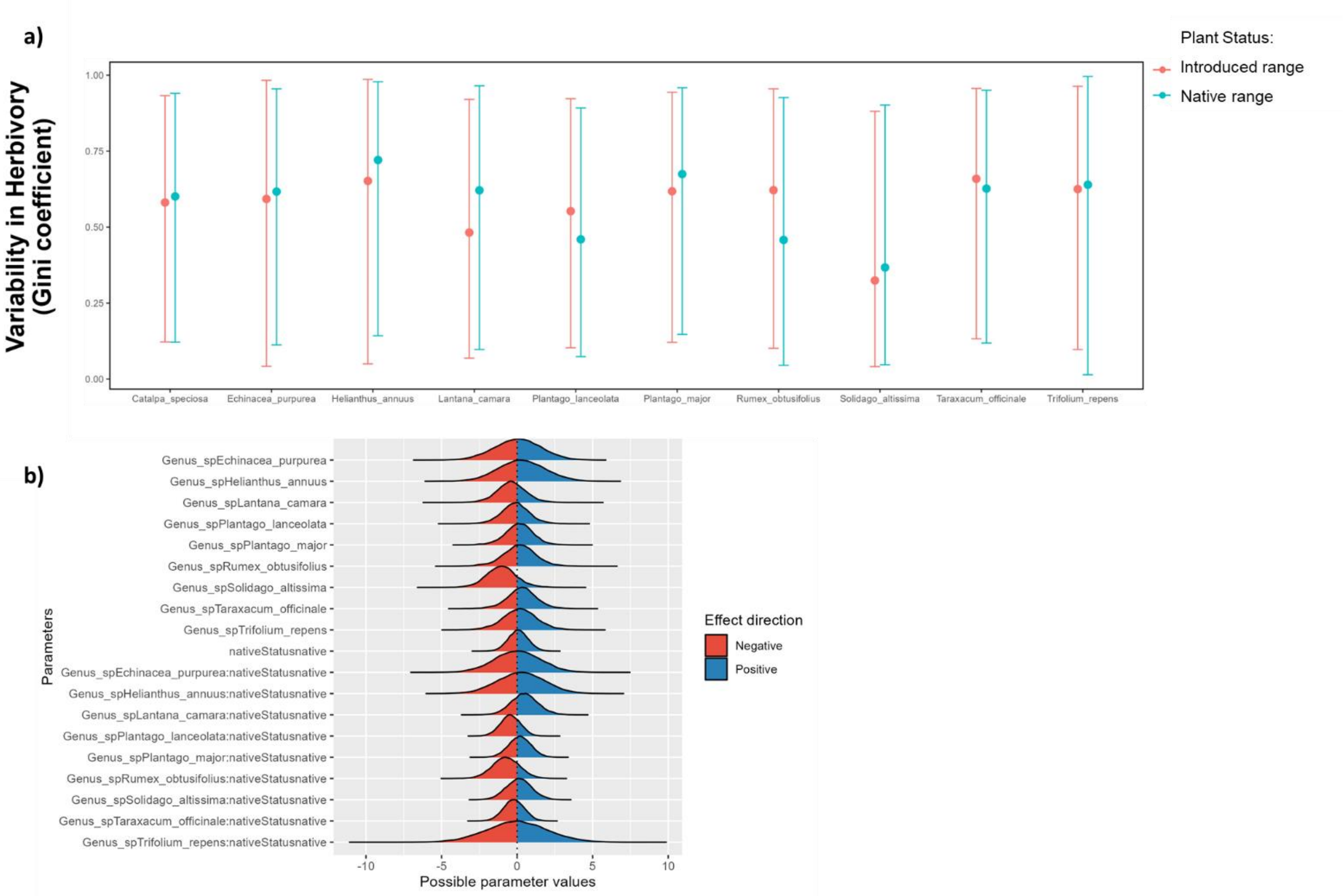


**Figure S2:** Results from models for the biogeographical subsets of ten species for variability in herbivory (Gini coefficient). a) Comparison of mean intrapopulation variability in herbivory for populations in their introduced (in red) and native (in blue) range for the subset of 10 species included in the biogeographic analysis and b) evaluation plot with the proportion of the posterior distribution of the factors in the model.

a)

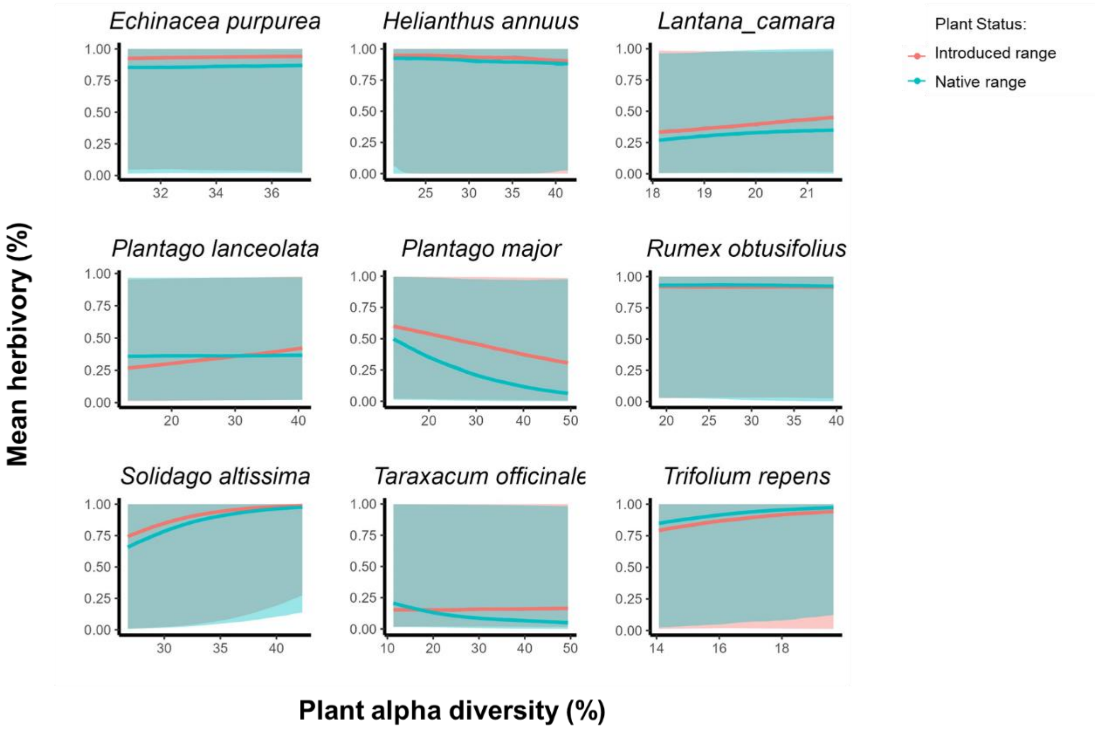


b)

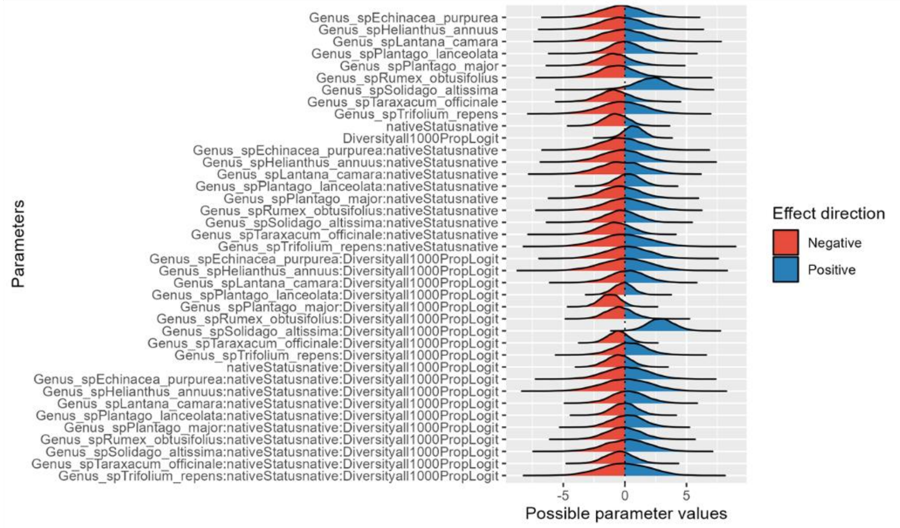


**Figure S3:** Results from models for the biogeographical subsets of nine species for the interaction between mean herbivory and plant diversity. a) Relationship between mean herbivory and surrounding plant alpha diversity for populations in their introduced (in red) and native (in blue) range for the subset of nine species included in the biogeographic analysis and b) evaluation plot with the proportion of the posterior distribution.

a)

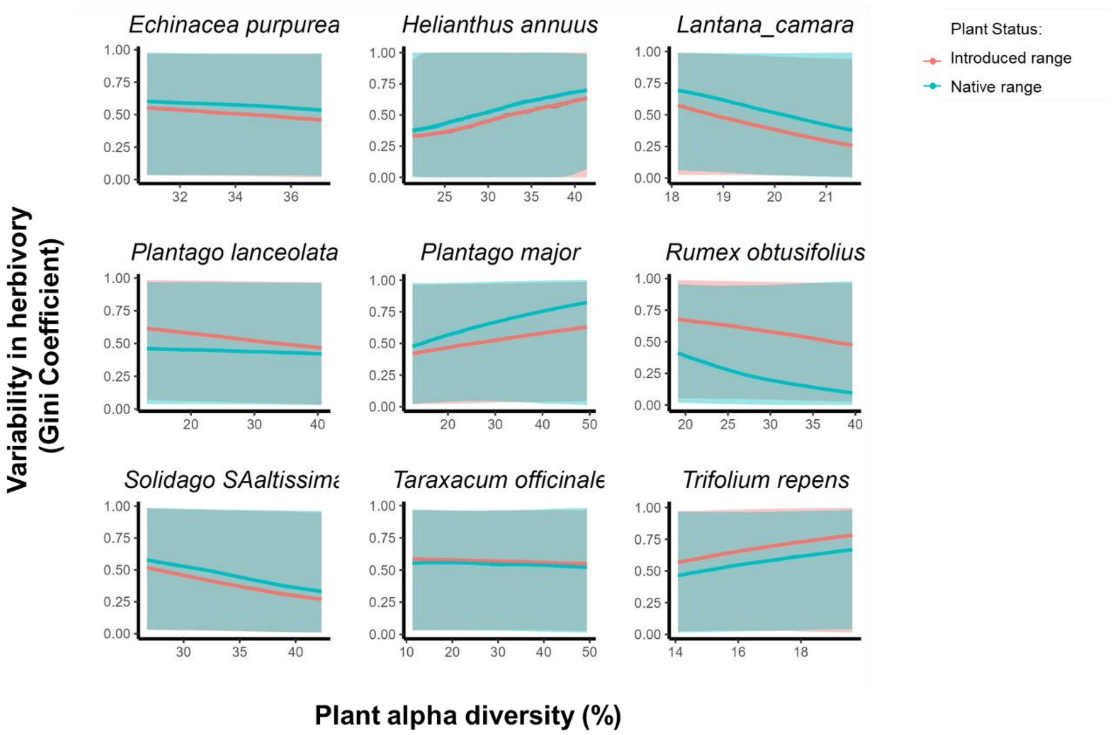


b)

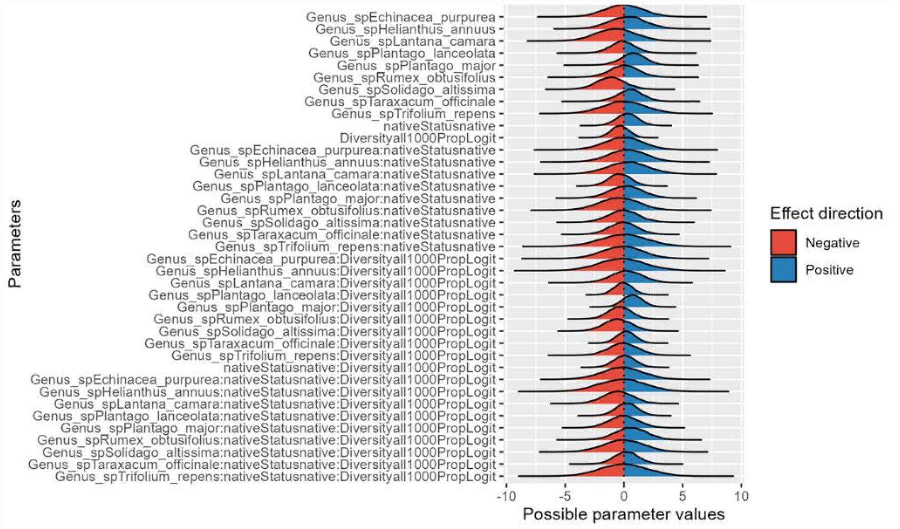


**Figure S4:** Results from models for the biogeographical subsets of nine species for the interaction between variability in herbivory (Gini coefficient) and plant diversity. a) Relationship between variability in herbivory and surrounding plant alpha diversity for populations in their introduced (in red) and native (in blue) range for the subset of nine species included in the biogeographic analysis and b) evaluation plot with the proportion of the posterior distribution.

a)

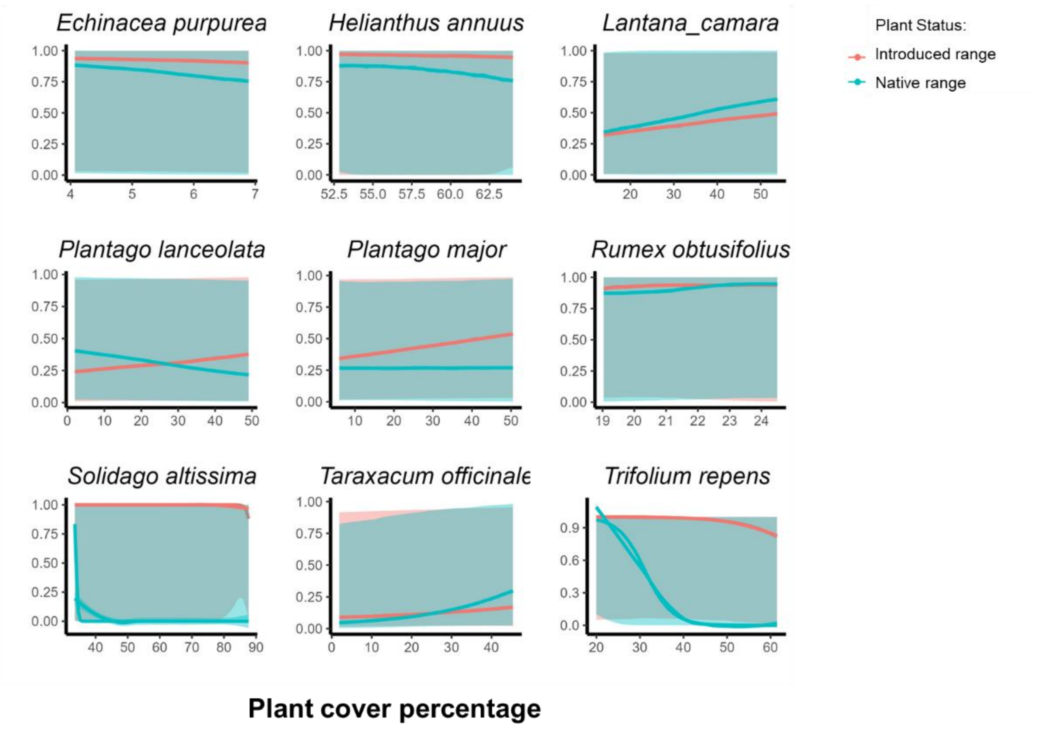


b)

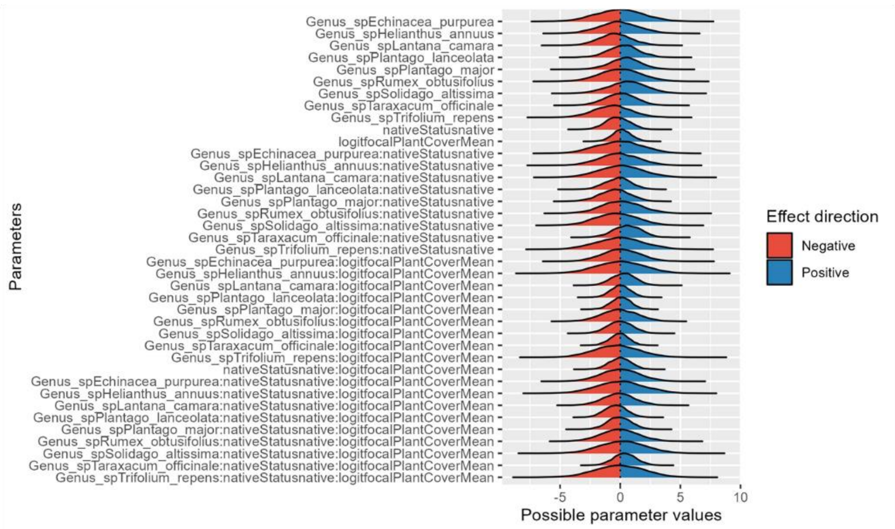


**Figure S5:** Results from models for the biogeographical subsets of ten species for the interaction between variability in herbivory (Gini coefficient) and plant cover percentage. a) Relationship between variability in herbivory and mean plant cover percentage for populations in their introduced (in red) and native (in blue) range for the subset of nine species included in the biogeographic analysis and b) evaluation plot with the proportion of the posterior distribution.

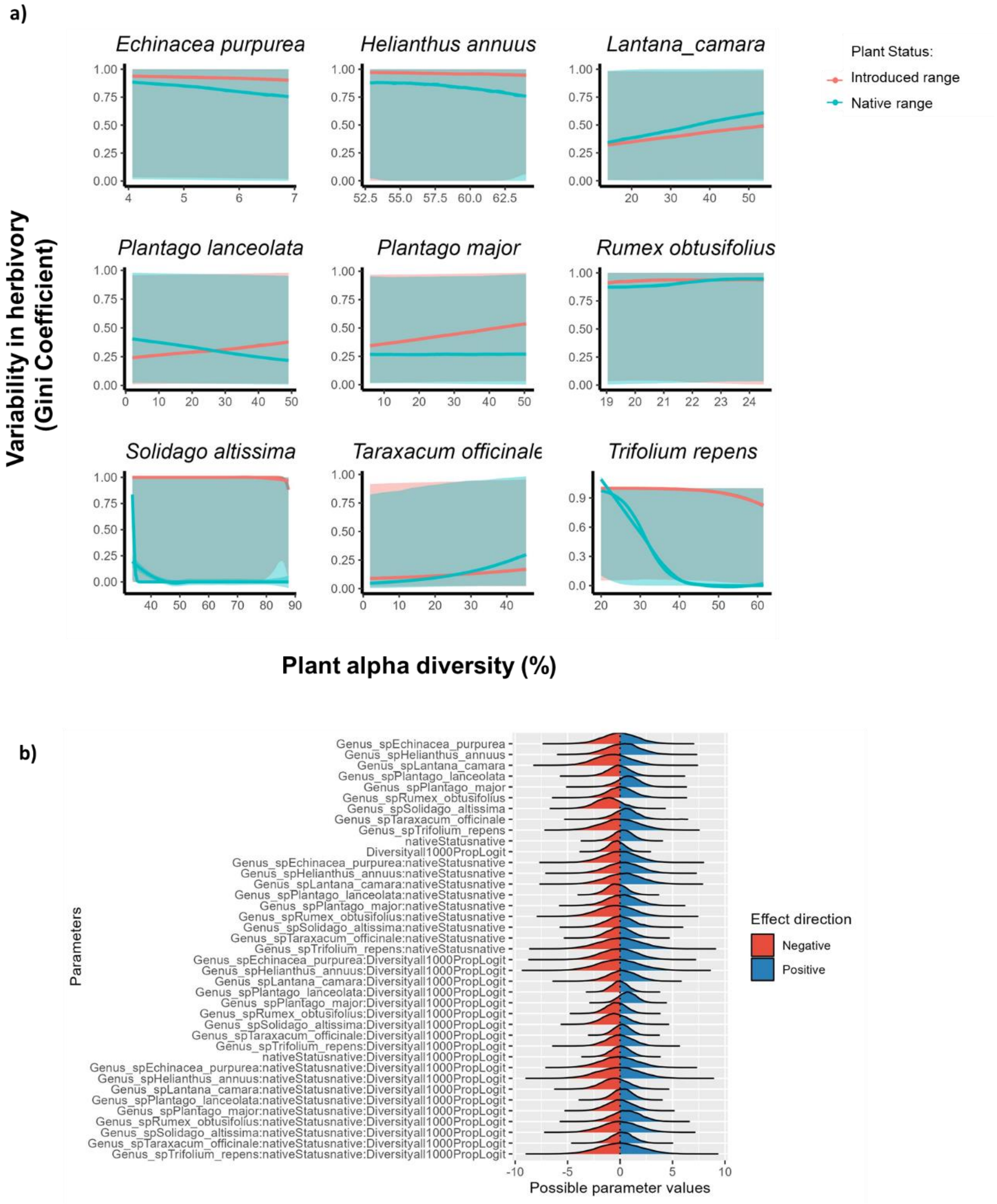


**Figure S6:** Results from models for the biogeographical subsets of ten species for the interaction between variability in herbivory (Gini coefficient) and plant cover percentage. a) Relationship between variability in herbivory and mean plant cover percentage for populations in their introduced (in red) and native (in blue) range for the subset of nine species included in the biogeographic analysis and b) evaluation plot with the proportion of the posterior distribution.

**Table S1**: Non-native species included in the study, indicating their invasion status (alien or naturalized) in their introduced range country and the number of populations per country.

| Species name | GloNAF status | Country | n Populations |
|---|---|---|---|
| *Abutilon theophrasti* | naturalized | United_States | 1 |
| *Acer campestre* | naturalized | United_States | 1 |
| *Achillea ptarmica* | naturalized | Finland | 1 |
| *Ageratum conyzoides* | naturalized | India | 1 |
| *Ailanthus altissima* | naturalized | United_States | 1 |
| *Alcea rosea* | naturalized | United_States | 1 |
| *Alliaria petiolata* | naturalized | United_States | 1 |
| *Alternanthera ficoidea* | naturalized | India | 1 |
| *Alternanthera sessilis* | alien | Nigeria | 2 |
| *Andropogon virginicus* | naturalized | Japan | 2 |
| *Arctium minus* | naturalized | United_States | 2 |
| *Argemone mexicana* | naturalized | India | 1 |
| *Asclepias curassavica* | naturalized | United_States | 1 |
| *Barbarea vulgaris* | naturalized | United_States | 1 |
| *Brassica oleracea* | naturalized | United_States | 1 |
| *Calotropis procera* | naturalized | South_Africa | 1 |
| *Campanula persicifolia* | naturalized | United_States | 1 |
| *Canna indica* | naturalized | South_Africa | 2 |
| *Castanea sativa* | naturalized | Spain | 4 |
| *Catalpa speciosa* | naturalized | United_States | 2 |
| *Centaurea stoebe* | naturalized | United_States | 1 |
| *Chenopodium album* | naturalized | United States | 1 |
| *Chromolaena odorata* | alien | Nigeria | 4 |
| *Crotalaria pallida* | naturalized | Brazil | 1 |
| *Ginkgo biloba* | naturalized | United_States | 1 |
| *Glycine max* | naturalized | United_States | 1 |
| *Gossypium hirsutum* | naturalized | Mexico | 2 |
| *Helianthus annuus* | naturalized | United States | 1 |
| *Heliotropium indicum* | alien | Mozambique | 1 |
| *Heterotheca subaxillaris* | naturalized | Israel | 1 |
| *Hypochaeris radicata* | naturalized | Australia | 1 |
| *Hypochaeris radicata* | naturalized | Japan | 3 |
| *Hyptis suaveolens* | naturalized | India | 1 |
| *Impatiens parviflora* | naturalized | United_Kingdom | 1 |
| *Lantana camara* | naturalized | India | 2 |
| *Lantana camara* | naturalized | Portugal | 2 |
| *Lepidium draba* | naturalized | United_States | 8 |
| *Lupinus polyphyllus* | naturalized | Finland | 1 |
| *Lythrum salicaria* | naturalized | United_States | 1 |
| *Melanthera scandens* | alien | Nigeria | 1 |
| *Morus alba* | naturalized | United_States | 1 |
| *Nandina domestica* | naturalized | United_States | 1 |

| | | | |
|---|---|---|---|
| *Oenothera biennis* | naturalized | Japan | 3 |
| *Olea europaea* | naturalized | New_Zealand | 1 |
| *Pastinaca sativa* | naturalized | United_States | 1 |
| *Plantago lanceolata* | naturalized | Australia | 1 |
| *Plantago lanceolata* | alien | Ecuador | 1 |
| *Plantago lanceolata* | naturalized | Japan | 12 |
| *Plantago lanceolata* | naturalized | New_Zealand | 8 |
| *Plantago lanceolata* | naturalized | Portugal | 2 |
| *Plantago lanceolata* | naturalized | Sweden | 3 |
| *Plantago lanceolata* | naturalized | United States | 15 |
| *Plantago major* | naturalized | Argentina | 1 |
| *Plantago major* | naturalized | China | 1 |
| *Plantago major* | alien | Ecuador | 1 |
| *Plantago major* | naturalized | New_Zealand | 2 |
| *Plantago major* | naturalized | United_States | 6 |
| *Potentilla recta* | naturalized | United_States | 1 |
| *Ruellia tuberosa* | naturalized | India | 1 |
| *Rumex crispus* | naturalized | United_States | 2 |
| *Salpichroa origanifolia* | naturalized | Portugal | 2 |
| *Senecio elegans* | naturalized | New_Zealand | 2 |
| *Senecio madagascariensis* | naturalized | Colombia | 3 |
| *Senecio viscosus* | naturalized | Finland | 1 |
| *Senna occidentalis* | naturalized | India | 1 |
| *Solanum dulcamara* | naturalized | United_States | 1 |
| *Solidago altissima* | naturalized | Japan | 8 |
| *Symphyotrichum novi-belgii* | naturalized | Japan | 2 |
| *Taraxacum campylodes* | naturalized | Argentina | 4 |
| *Taraxacum campylodes* | naturalized | Australia | 1 |
| *Taraxacum campylodes* | naturalized | Colombia | 3 |
| *Taraxacum campylodes* | alien | Ecuador | 1 |
| *Taraxacum campylodes* | naturalized | Japan | 3 |
| *Taraxacum campylodes* | naturalized | Mexico | 2 |
| *Taraxacum campylodes* | naturalized | New_Zealand | 4 |
| *Taraxacum campylodes* | naturalized | United_States | 8 |
| *Tradescantia fluminensis* | naturalized | Portugal | 2 |
| *Tragopogon dubius* | naturalized | United States | 1 |
| *Trifolium repens* | naturalized | Portugal | 2 |
| *Urtica dioica* | naturalized | United States | 1 |
| *Verbascum thapsus* | naturalized | United_States | 1 |
| *Verbesina encelioides* | naturalized | Israel | 1 |
| *Vicia sativa* | naturalized | United_States | 1 |
| *Vicia villosa* | naturalized | United_States | 1 |
| *Zea mays* | naturalized | United_States | 1 |

**Table S2:** subset of ten species in their native and introduced ranges indicating the occurrence country and the number of populations on each range and country.

| Species name | Country | Plant status | n Populations |
|---|---|---|---|
| *Catalpa_speciosa* | United States | introduced | 3 |
| *Catalpa_speciosa* | United States | native | 1 |
| *Echinacea_purpurea* | United States | introduced | 4 |
| *Echinacea_purpurea* | United States | native | 2 |
| *Helianthus_annuus* | United States | introduced | 1 |
| *Helianthus_annuus* | United States | native | 1 |
| *Lantana_camara* | Colombia | native | 1 |
| *Lantana_camara* | India | introduced | 2 |
| *Lantana_camara* | Portugal | introduced | 1 |
| *Plantago_lanceolata* | Australia | introduced | 1 |
| *Plantago_lanceolata* | Canada | introduced | 1 |
| *Plantago_lanceolata* | China | native | 1 |
| *Plantago_lanceolata* | Ecuador | introduced | 1 |
| *Plantago_lanceolata* | Finland | native | 1 |
| *Plantago_lanceolata* | Germany | native | 4 |
| *Plantago_lanceolata* | Ireland | native | 3 |
| *Plantago_lanceolata* | Japan | introduced | 2 |
| *Plantago_lanceolata* | New Zealand | introduced | 4 |
| *Plantago_lanceolata* | Portugal | introduced | 1 |
| *Plantago_lanceolata* | Spain | native | 8 |
| *Plantago_lanceolata* | Sweden | native | 4 |
| *Plantago_lanceolata* | Switzerland | native | 12 |
| *Plantago_lanceolata* | United Kingdom | native | 1 |
| *Plantago_lanceolata* | United States | introduced | 15 |
| *Plantago_lanceolata* | United States | native | 1 |
| *Plantago_major* | Argentina | introduced | 1 |
| *Plantago_major* | China | native | 1 |
| *Plantago_major* | Ecuador | introduced | 1 |
| *Plantago_major* | Estonia | native | 1 |
| *Plantago_major* | Finland | native | 2 |
| *Plantago_major* | India | introduced | 3 |
| *Plantago_major* | New Zealand | introduced | 1 |
| *Plantago_major* | Panama | introduced | 1 |
| *Plantago_major* | United States | introduced | 8 |
| *Rumex_obtusifolius* | Canada | introduced | 1 |
| *Rumex_obtusifolius* | Panama | introduced | 1 |
| *Rumex_obtusifolius* | Spain | native | 2 |
| *Solidago_altissima* | Japan | introduced | 3 |
| *Solidago_altissima* | United States | native | 2 |
| *Taraxacum_officinale* | Argentina | introduced | 4 |
| *Taraxacum_officinale* | Australia | introduced | 3 |
| *Taraxacum_officinale* | Chile | introduced | 2 |

| | | | |
|---|---|---|---|
| *Taraxacum_officinale* | Colombia | introduced | 1 |
| *Taraxacum_officinale* | Ecuador | introduced | 1 |
| *Taraxacum_officinale* | Finland | native | 2 |
| *Taraxacum_officinale* | Germany | native | 7 |
| *Taraxacum_officinale* | Japan | introduced | 1 |
| *Taraxacum_officinale* | Mexico | introduced | 1 |
| *Taraxacum_officinale* | New Zealand | introduced | 2 |
| *Taraxacum_officinale* | Norway | native | 1 |
| *Taraxacum_officinale* | Spain | native | 1 |
| *Taraxacum_officinale* | Switzerland | native | 2 |
| *Taraxacum_officinale* | United Kingdom | native | 1 |
| *Taraxacum_officinale* | United States | introduced | 8 |
| *Trifolium_repens* | Germany | native | 1 |
| *Trifolium_repens* | Portugal | introduced | 1 |
| *Trifolium_repens* | United Kingdom | native | 1 |
